# Species and Accession Diversity of Secondary Metabolites and Antioxidant Activity in Legume Sprouts

**DOI:** 10.64898/2026.07.31.741982

**Authors:** Gyeongseon An, Hakyung Kwon, Jungmin Ha

## Abstract

Legume sprouts contain diverse bioactive phytochemicals, yet species- and accession-level comparisons of antioxidant properties and secondary metabolites remain limited. Antioxidant capacity, total phenolic content (TPC), total flavonoid content (TFC), and 19 secondary metabolites were evaluated in sprouts of soybean (*Glycine max* (L.) Merr.), mungbean (*Vigna radiata* (L.) R. Wilczek), cowpea (*Vigna unguiculata* (L.) Walp.), and peanut (*Arachis hypogaea* L.), using ten accessions per species under standardized conditions. Mungbean and cowpea sprouts showed significantly higher antioxidant activity, TPC, and TFC than soybean and peanut; across accessions, ABTS and DPPH activities ranged from 13.67 to 49.33% and 7.91 to 55.16%, and TPC and TFC from 3.91 to 13.81 mg GAE/g and 0.05 to 1.04 mg QE/g, respectively. Metabolite profiling revealed species-specific patterns, including isoflavones in soybean, rutin in mungbean, coumestrol in cowpea and resveratrol in peanut. Both antioxidant and phytochemical profiles varied with species and accession, so both can be selected to obtain sprouts with targeted properties.

## 1. introduction

Legumes are an important source of protein in developing countries and contain essential minerals necessary for human health.^1–2^ In addition to their nutritional value, legumes synthesize a wide range of secondary metabolites to protect themselves from biotic and abiotic stresses, such as pest attack and ultraviolet radiation. Some secondary metabolites found in legumes can contribute to human health, and their types and concentrations show variations among species.

Vitexin, which is abundant in mungbean (*Vigna radiata* (L.) R. Wilczek), has been reported to exhibit strong antioxidant and neuroprotective effects.^3^ Similarly, resveratrol, a major phenolic secondary metabolite in peanut (*Arachis hypogaea* L.), may help reduce the risk of Alzheimer’s disease.^4^ Soybean (*Glycine max* (L.) Merr.), a representative legume crop, is particularly rich in isoflavones, which are legume-specific secondary metabolites.^5^ Isoflavones are flavonoid secondary metabolites structurally similar to estrogen, a female steroid hormone, and have been shown to provide various health benefits, including alleviation of menopausal hot flashes and prevention of osteoporosis.^6–8^ As various secondary metabolites with potential health benefits have been identified in legumes, numerous studies have been conducted to increase the concentrations of these secondary metabolites in legumes.^9–10^

Germination is a method that can change the concentrations of secondary metabolites by activating related biosynthetic enzymes with relatively low costs.^11–12^ In soybean, isoflavone concentrations increase during germination.^13^ Likewise, germination enhances the levels of phenolic acids and flavonoids—such as gallic acid, caffeic acid, p-coumaric acid, and quercetin—in mungbean. Notably, while chlorogenic acid was not detected in mungbean seeds, it was identified in mungbean sprouts.^14^ Regarding pea sprouts, the concentrations of gallic acid, catechin, syringic acid, and quercetin increased significantly during germination. Furthermore, vanillin and ferulic acid, which were absent in seeds, were newly detected upon sprouting.^15^

Accordingly, because germination of legumes can alter secondary metabolite profiles, this process has been widely utilized in studies aimed at enhancing their functional properties. However, while the concentrations of individual secondary metabolites have been reported for various legume sprouts, few studies have profiled these metabolites across multiple species and diverse accessions under identical environmental conditions.

Therefore, this study aimed to compare the secondary metabolite profiles both within and between species, using ten accessions each of soybean (*Glycine max* (L.) Merr.), mungbean (*Vigna radiata* (L.) R. Wilczek), cowpea (*Vigna unguiculata* (L.) Walp.), and peanut (*Arachis hypogaea* L.) sprouts, all cultivated under standardized environmental conditions. Antioxidant activity, total phenolic content (TPC), and total flavonoid content (TFC) were also measured to evaluate the functional properties of each legume sprout. These findings are expected to provide fundamental data for cultivar improvement and selective utilization of legume sprouts with enhanced functional properties.

## 2. Materials and methods

### 2.1 Sample preparation

Soybean samples were harvested from the experimental farm of Seoul National University, Suwon, Republic of Korea (37°16′12.094″ N, 126°59′20.756″ E). Mungbean, cowpea, and peanut seeds were provided by Kangwon National Universitye, Chonnam National University, and Pusan National University in Korea, respectively (Figure 1). Ten accessions per species were used in this study (Table S1).^16^ Seeds were washed and then immersed in dark conditions at 37°C for 6 hours in an incubator (JEIO TECH, Daejeon, Korea). The seeds that had been immersed were then germinated for 4 days under irrigation conditions of 2 minutes every 4 hours at 28 ± 2°C using a sprout cultivation machine (Sundotcom, Gongju, Korea).^17^ Soybean, mungbean, cowpea, and peanut sprouts grown for 4 days were dried at 70°C for 24 hours and then pulverized. The resulting sprout powder was extracted with 70% ethanol for 24 hours and centrifuged at 3000 rpm for 15 minutes, and the supernatant was filtered through a 0.4 μm syringe filter. The filtered sprout extract was stored at 4°C until further experiments.

**Figure 1.**
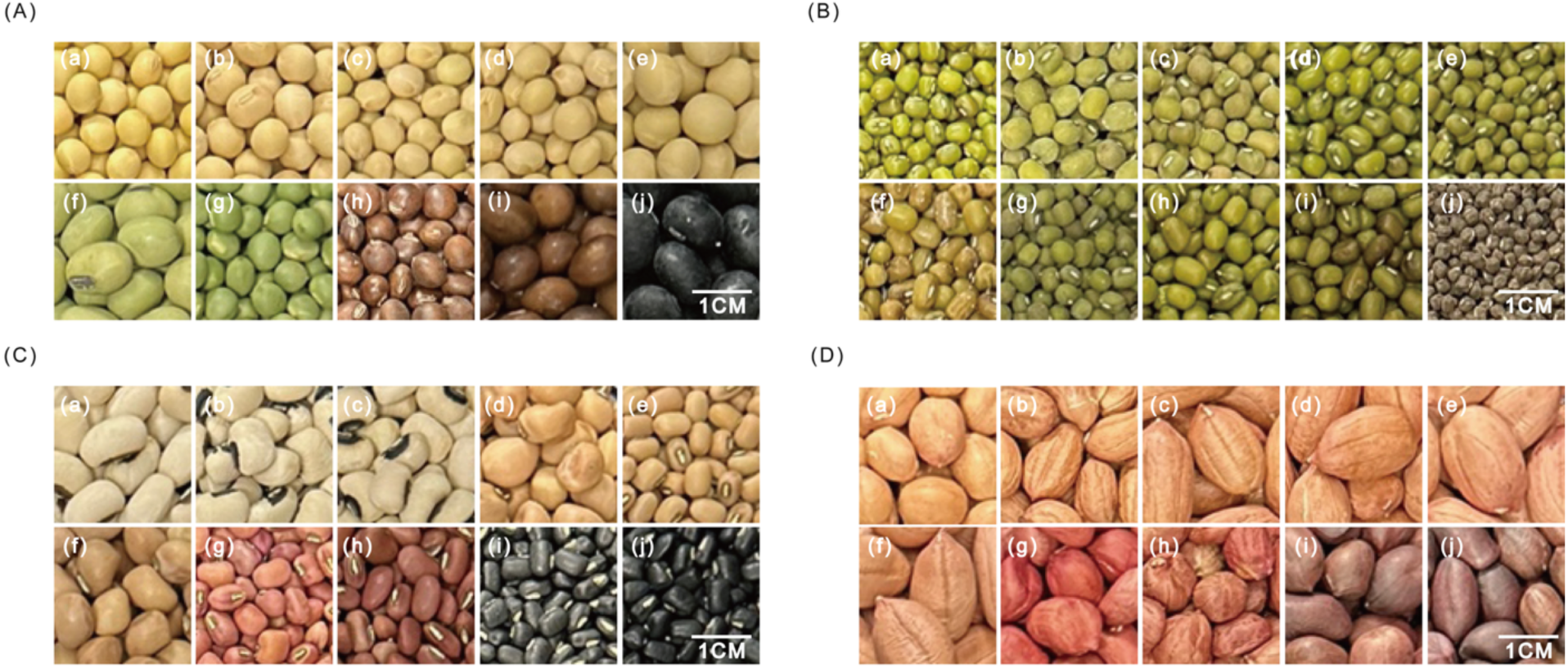
Phenotypes of soybean, mungbean, cowpea, and peanut seeds used in this study. **(A)**; Haepum (a), Joyang (b), SG188 (c), Pungsankong (d), Daechan (e), Cheongminin (f), Gilyuk102 (g), SG151 (h), Chamame (i), Seoritae (j) **(B)**; M357 (a), M369 (b), Sanpo (c), M658 (d), M243 (e), M366 (f), M625 (g), Seonhwa (h), M334 (i), M730 (j) **(C);** C365 (a), C319 (b), C292 (c), C378 (d), C242( e), C369 (f), C282 (g), C123 (h), C81 (i), C93 (j) **(D)**; P189 (a), P111 (b), P319 (c), P177 (d), P277 (e), P294 (f), P310 (g), P378 (h), P191 (i), P366 (j)

### 2.2 Antioxidant activity measurement

#### 2.2.1 2,2’-azino-bis(3-ethylbenzothiazoline-6-sulfonic acid) (ABTS) radical scavenging activity

ABTS radical scavenging activity was performed by slightly modifying the method proposed by Pasri *et al*.^18^ ABTS 7.4 mM solution Sigma-Aldrich (St. Louis, MO, USA) and Potassium persulfate 2.6 mM solution (Yakuri, Kyoto, Japan) were mixed at a 1:1 ratio and stored under dark conditions for 24 hours. The mixed solution was diluted with phosphate-buffered saline and adjusted to have an absorbance value of 0.70 ± 0.03 at 740 nm. Ascorbic acid (Fujifilm Wako Pure Chemical Corporation, Osaka, Japan) was used as a standard at concentrations of 0, 10, 20, 40, and 60 mg/L. After adding 20 μL of each sample (1 g/L) and standard to a 96-well microplate, 180 μL of the diluted ABTS solution was added. After reacting in the dark for 10 minutes, the absorbance value was measured at 740 nm using a microplate reader (Thermo Fisher Scientific, Waltham, MA, USA).

#### 2.2.2 2,2-diphenyl-1-picrylhydrazyl (DPPH) radical scavenging activity

DPPH radical scavenging activity was measured using The OxiTec™ DPPH Antioxidant Assay Kit (BIOMAX, Gyeonggi, Korea). As a standard, a 6-hydroxy-2,5,7,8-tetramethylchroman-2-carboxylic acid (0, 10, 60 and 80 mg/L) was used. According to the DPPH radical scavenging activity measurement method suggested by the manufacturer, 20 μL sample (10 g/L) and standard were put in a 96-well microplate, and 80 μL of assay buffer and 100 μL of DPPH working solution were added. After reacting in the dark for 30 minutes, the absorbance value was measured at 520 nm with a microplate reader (Thermo Fisher Scientific, Waltham, MA, USA).

### 2.3 Determination of total phenolic and flavonoid concentrations

#### 2.3.1 Total flavonoid content

Total flavonoid content (TFC) was determined using the aluminum nitrate colorimetric method described by Chang et al. with slight modifications.^19^ A reaction solution was prepared by mixing 1 M potassium acetate (Fujifilm Wako Pure Chemical Corporation, Osaka, Japan) and 10% aluminum nitrate (Junsei Chemical Co., Ltd., Tokyo, Japan) at a 1:1 ratio. Quercetin Sigma-Aldrich (St. Louis, MO, USA) was used as a standard at concentrations of 0, 50, 100, and 200 mg/L. Each sample (50 g/L) or standard was mixed with the reaction solution and incubated for 30 min. Absorbance was measured at 405 nm using a microplate spectrophotometer (Thermo Fisher Scientific, Waltham, MA, USA).

#### 2.3.2 Total phenolic content

Total phenolic content (TPC) was determined using the Folin–Ciocalteu assay described by Singleton et al. with minor modifications.^20^ Gallic acid Sigma-Aldrich (St. Louis, MO, USA) was used as a standard at concentrations of 0, 50, 100, and 200 mg/L. A 100 μL aliquot of each sample or standard was mixed with 50 μL of Folin–Ciocalteu reagent Sigma-Aldrich (St. Louis, MO, USA) and incubated for 5 min. Then, 300 μL of 20% Na₂CO₃ solution (Fujifilm Wako Pure Chemical Corporation, Osaka, Japan) was added, and the mixture was allowed to react for 15 min under dark conditions. Finally, 1 mL of distilled water was added, and the absorbance of the supernatant obtained by centrifugation was measured at 740 nm using a microplate spectrophotometer (Thermo Fisher Scientific, Waltham, MA, USA).

### 2.4 Identification and quantification of individual metabolites

Individual flavonoids and phenolic acids were identified and quantified using a ZORBAX SB-C18 column (3.5 μm, 4.6 mm × 150 mm, Agilent Technologies, Santa Clara, CA, USA) on an ultra-performance liquid chromatography (UPLC, Shimadzu Corporation, Kyoto, Japan) system equipped with a photodiode array detector. Standard curves were established using the following standards purchased from ChemFaces (Wuhan, China): gallic acid, neochlorogenic acid, 4-hydroxybenzoic acid, caffeic acid, chlorogenic acid, vanillic acid, p-coumaric acid, trans-ferulic acid, cinnamic acid, daidzin, glycitin, rutin, vitexin, isoquercitrin, resveratrol, daidzein, genistein, coumestrol, and biochanin A. The mobile phase consisted of 0.1% acetic acid (solvent A) and acetonitrile (solvent B). A gradient elution program was applied as follows: 0–10 min, 95% A; 10–11 min, 95–90% A; 11–20 min, 90% A; 20.1–22 min, 80% A; 22–23 min, 80–90% A; 23–24 min, 90–95% A; 24–29 min, 95–65% A; 29–35 min, 65–50% A; 35.1–50 min, 95% A. The flow rate was 1 mL/min, injection volume was 2 μL, and the column oven temperature was maintained at 40°C. Detection wavelengths were set at 260 nm for 4-hydroxybenzoic acid, 340 nm for vitexin and isoquercitrin, 240 nm for coumestrol, and 280 nm for the other 15 standards (Table S2).

### 2.5 Statistical analysis

Statistical significance was evaluated by analysis of variance (ANOVA) followed by Duncan’s multiple range test (*p* < 0.05). Pearson correlation analysis was conducted among ABTS radical scavenging activity, DPPH radical scavenging activity, total flavonoid content, total phenolic content, and selected secondary metabolites in each species. The secondary metabolites included in the correlation analysis were selected based on their classification in the highest statistical group according to post hoc multiple comparison analysis. Chlorogenic acid exhibited the highest mean concentration in peanut sprouts among the four species. Nevertheless, several mungbean accessions displayed concentration ranges comparable to those of peanut sprouts. Therefore, chlorogenic acid was additionally included in the correlation analysis of mungbean sprouts.

All experiments were performed in triplicate, and statistical analyses were conducted using R software (version 4.2.0). Correlation coefficients were interpreted according to the guidelines described by Haldun Akoglu *et al*.^21^

## 3. Results

### 3.1 Measurement of total flavonoid and phenolic contents

Soybean, mungbean, cowpea, and peanut sprouts showed significant variations in TFC by species and by accessions within species (Figure 2A and C). The ranges of TFC for each species were 0.14 mg QE/g ± 0.01 mg QE/g to 0.35 mg QE/g ± 0.01 mg QE/g for soybean sprouts, 0.52 mg QE/g ± 0.02 mg QE/g to 0.82 mg QE/g ± 0.02 mg QE/g for mungbean sprouts, 0.47 mg QE/g ± 0.02 mg QE/g to 1.04 mg QE/g ± 0.02 mg QE/g for cowpea sprouts, and 0.05 mg QE/g ± 0.00 mg QE/g to 0.29 mg QE/g ± 0.01 mg QE/g for peanut sprouts (Figure 2A). Among the four species, TFC was significantly higher in mungbean and cowpea sprouts than in soybean and peanut sprouts. For soybean sprouts, Daechan (0.35 mg QE/g ± 0.01 mg QE/g) and Pungsankong (0.32 mg QE/g ± 0.05 mg QE/g) exhibited the highest TFC, M658 (0.82 mg QE/g ± 0.02 mg QE/g) for mungbean sprouts, C93 (1.02 mg QE/g ± 0.03 mg QE/g), C242 (1.04 mg QE/g ± 0.03 mg QE/g) for cowpea sprouts and P191 (0.29 mg QE/g ± 0.01 mg QE/g) for peanut sprouts had the highest TFC (Figure 2C). TPC, like TFC, also showed diverse variations among species and accessions (Figures 2B and 2D). The range of TPC by species was 3.91 mg GAE/g ± 0.01 mg GAE/g to 7.77 mg GAE/g ± 0.06 mg GAE/g for soybean sprouts, 11.50 mg GAE/g ± 0.03 mg GAE/g to 13.81 mg GAE/g ± 0.06 mg GAE/g for mungbean sprouts, 7.96 mg GAE/g ± 0.03 mg GAE/g to 13.80 mg GAE/g ± 0.06 mg GAE/g for cowpea sprouts, and 4.10 mg GAE/g ± 0.15 mg GAE/g to 6.64 mg GAE/g ± 0.50 mg GAE/g for peanut sprouts (Figures 2B). The TPC of mungbean and cowpea sprouts was significantly higher than those of soybean and peanut sprouts, and there was no significant difference between soybean and peanut sprouts. Unlike TFC, TPC in mungbean sprouts was significantly higher than TPC in cowpea sprouts. Daechan (7.77 mg GAE/g ± 0.06 mg GAE/g) in soybean sprouts, M243 (13.81 mg GAE/g ± 0.06 mg GAE/g) in mungbean sprouts, C81 (11.80 mg GAE/g± 0.03 mg GAE/g) in cowpea sprouts, and P277 (6.64 mg GAE/g ± 0.50 mg GAE/g) in peanut sprouts showed the highest TPC (Figure 2D).

**Figure 2.**
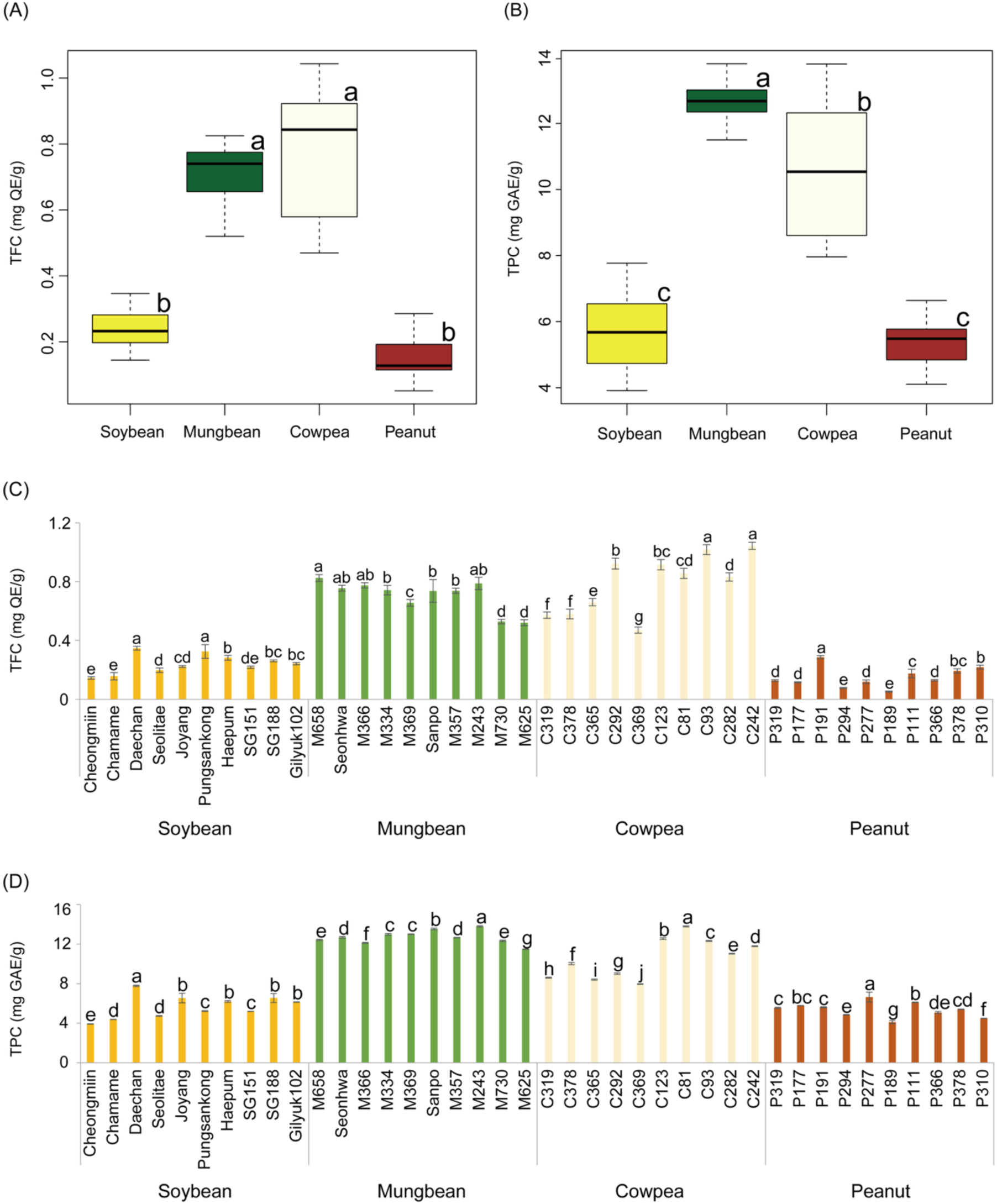
Total flavonoid content (TFC) and total phenolic content (TPC) of soybean, mungbean, cowpea, and peanut sprouts Figures 2A and 2B present interspecific variation in TFC and TPC, respectively, among soybean, mungbean, cowpea, and peanut sprouts. Figures 2C and 2D present intraspecific variation in TFC and TPC, respectively. Different lowercase letters indicate significant differences among species (A and B) or among accessions within each species (C and D), according to Duncan’s multiple range test (p < 0.05).

### 3.2 Antioxidant activity

Antioxidant activities of soybean, mungbean, cowpea, and peanut sprouts were evaluated by ABTS and DPPH radical scavenging assays (Figure 3). The range of ABTS radical scavenging activity was 13.67% ± 0.01% to 31.65% ± 0.01% for soybean sprouts, 35.92% ± 0.01% to 44.96% ± 0.01% for mungbean sprouts, 26.05% ± 0.01% to 49.33% ± 0.01% for cowpea sprouts, and 14.34% ± 0.00% to 22.35% ± 0.01% for peanut sprouts (Figure 3A). The range of DPPH radical scavenging activity was 16.35% ± 2.24% to 31.03% ± 1.00% for soybean sprouts, 34.65% ± 2.14% to 43.23% ± 0.84% for mungbean sprouts, 28.75% ± 0.19% to 55.16% ± 1.87% for cowpea sprouts, and 7.91% ± 1.43% to 22.45% ± 6.26% for peanut sprouts (Figure 3B). These results indicate that mungbean and cowpea sprouts exhibit significantly higher antioxidant activity than soybean and peanut sprouts, while peanut sprouts showed the lowest activity.

**Figure 3.**
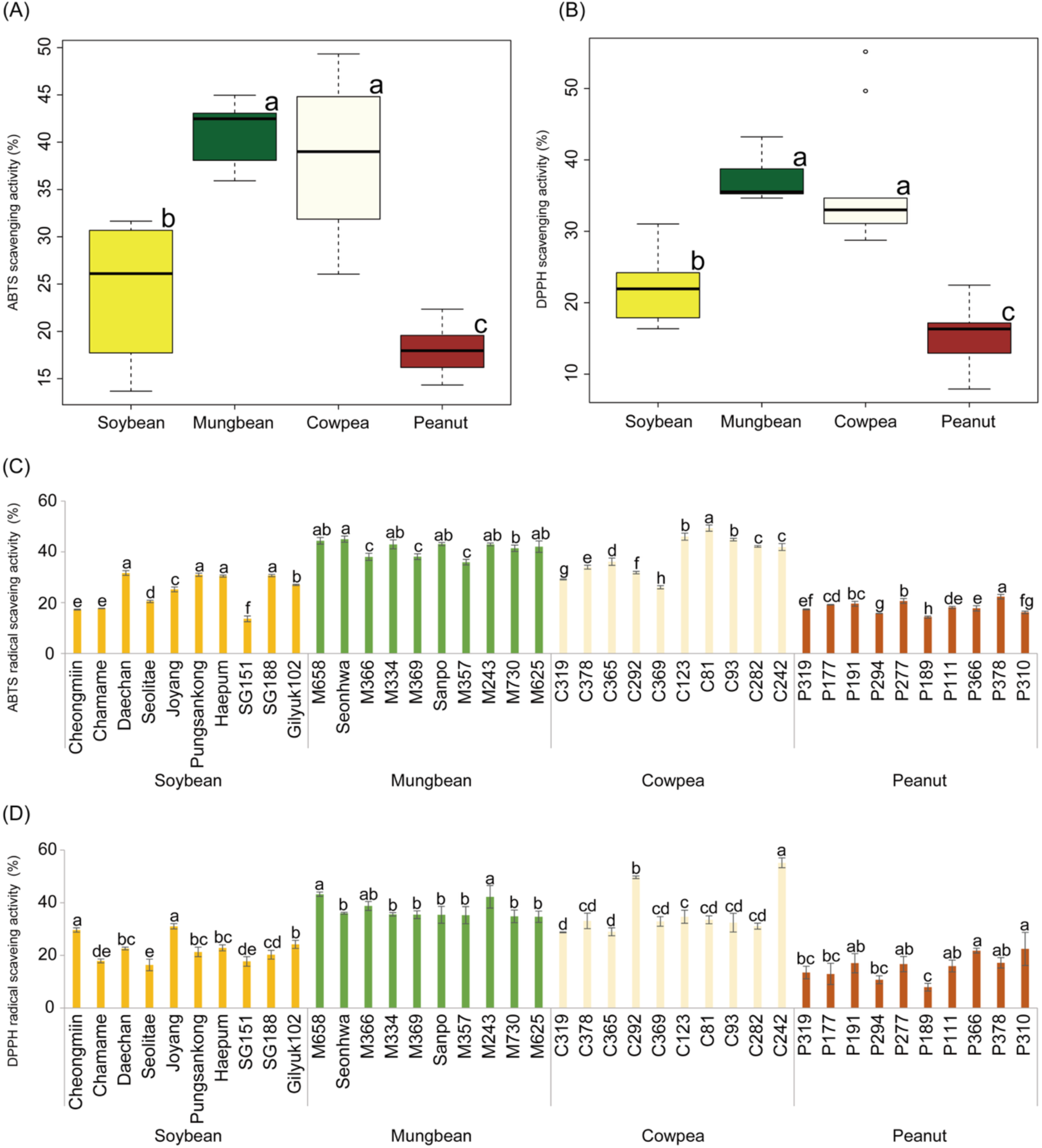
Comparison of antioxidant activities of soybean, mungbean, cowpea, and peanut sprouts. Figures 3A and 3B present interspecific ABTS radical scavenging activity (A) and DPPH radical scavenging activity (B), respectively, among soybean, mungbean, cowpea, and peanut sprouts. Figures 3C and 3D present intraspecific ABTS radical scavenging activity (C) and DPPH radical scavenging activity (D). Different lowercase letters indicate significant differences among species (A and B) or among accessions within each species (C and D), according to Duncan’s multiple range test (*p* < 0.05).

Within species comparisons revealed that in soybean sprouts, Daechan (31.65% ± 0.01%), Pungsankong (30.99% ± 0.01%), Haepum (30.47% ± 0.00%), and SG188 (30.68% ± 0.00%) exhibited significantly higher ABTS radical scavenging activity than the other soybean accessions (Figure 3C). Similarly, Seonhwa showed the highest ABTS activity among the ten mungbean accessions (44.96% ± 0.01%), C81 among the ten cowpea accessions (49.33% ± 0.01%), and P378 among the ten peanut accessions (22.35% ± 0.01%). For DPPH radical scavenging activity, Cheongmiin (29.62% ± 0.81%) and Joyang (31.03% ± 1.00%) showed the highest activity among the ten soybean accessions, M658 (43.23% ± 0.84%) and M243 (42.23% ± 4.29%) among the ten mungbean accessions, C242 (55.16% ± 1.87%) among the ten cowpea accessions, and P310 (22.45% ± 6.26%) among the ten peanut accessions (Figure 3D).

### 3.3 Quantification of Individual secondary metabolites in various legume sprouts

The concentrations of Individual secondary metabolite concentrations of 10 accessions per species were measured using UPLC (Figure 4). A total of 19 secondary metabolites were qualified and quantified. The secondary metabolite concentration showed significant variation within each species. The highest phenolic acid concentrations identified in each species were 8.13 ± 0.05 mg/L of chlorogenic acid in Haepum soybean sprouts, 13.84 ± 0.11 mg/L of chlorogenic acid in M369 mungbean sprouts, 9.06 ± 0.06 mg/L of gallic acid in C123 cowpea sprouts, and 11.68 ± 0.02 mg/L of chlorogenic acid in P319 peanut sprouts. The highest flavonoid concentrations within each species were 97.36 ± 0.45 mg/L of daidzin in Daechan soybean sprouts, 50.99 ± 0.24 mg/L of rutin in M334 mungbean sprouts, 53.58 ± 0.14 mg/L of daidzin in C242 cowpea sprouts, and 8.12 ± 0.02 mg/L of daidzein in P111 peanut sprouts.

**Figure 4.**
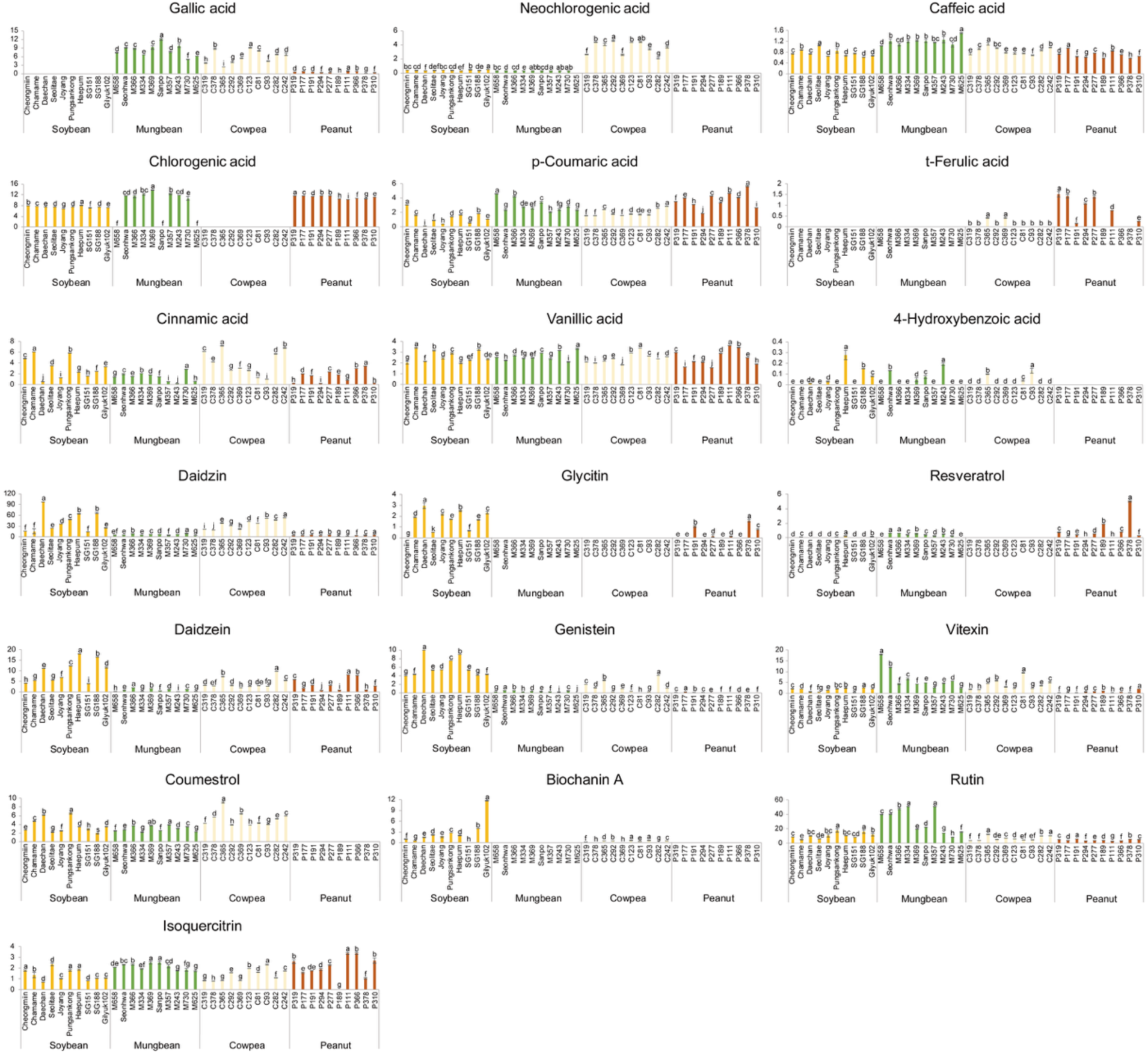
Concentrations of individual secondary metabolites in soybean, mungbean, cowpea, and peanut sprouts Individual secondary metabolite content of soybean, mungbean, cowpea, and peanut sprouts. Each lowercase letter indicates statistically significant differences based on intraspecies comparisons (*p* < 0.05).

Significant inter-species variations in secondary metabolite profiles were observed. For phenolic acids, the highest gallic acid concentration was observed in mungbean sprouts (5.09 ± 0.13 to 11.96 ± 0.32 mg/L), followed by cowpea sprouts (2.30 ± 0.03 to 9.06 ± 0.06 mg/L) and peanut sprouts (0.08 ± 0.00 to 0.97 ± 0.02 mg/L). In soybean sprouts, gallic acid was not detected in any of the ten accessions. Cowpea sprouts showed significantly higher amounts of neochlorogenic acid, ranging from 2.06 mg/L ± 0.07 mg/L to 4.57 mg/L ± 0.12 mg/L, whereas only trace amounts or no detectable levels were observed in soybean, mungbean, and peanut sprouts. The concentration of caffeic acid in mungbean sprouts ranged from 1.03 mg/L ± 0.00 mg/L to 1.50 mg/L ± 0.02 mg/L, which was significantly higher than that of soybean, cowpea, and peanut sprouts. In the case of t-ferulic acid, it was not detected in soybean and mungbean sprouts, while its concentration in cowpea sprouts ranged from not detected to 0.41 ± 0.01 mg/L. On the other hand, the t-Ferulic acid in peanut sprouts was detected upto 1.51 mg/L ± 0.13 mg/L, which was significantly higher than that in soybean, mungbean, and cowpea sprouts.

Among the flavonoids, glycitin, daidzein, genistein, and biochanin A in soybean sprouts ranged from 0.00 ± 0.00 to 4.40 ± 0.03 mg/L, 2.82 ± 0.05 to 18.02 ± 0.04 mg/L, 3.45 ± 0.03 to 9.96 ± 0.01 mg/L, and 0.00 ± 0.00 to 11.75 ± 0.11 mg/L, respectively, and these concentrations were significantly higher than those observed in mungbean, cowpea, and peanut sprouts. In contrast, mungbean sprouts contained higher amounts of vitexin (2.36 ± 0.00 to 17.90 ± 0.06 mg/L) and rutin (8.28 ± 0.21 to 50.99 ± 0.24 mg/L) than the other three species. Cowpea sprouts also showed higher coumestrol levels (3.81 ± 0.03 to 9.07 ± 0.10 mg/L). Resveratrol, a stilbene-type secondary metabolite, was detected at higher levels in peanut sprouts than in the other species (0.00 ± 0.00 to 5.04 ± 0.08 mg/L).

### 3.4 Correlation analysis between antioxidant activity, bioactive substances and Individual secondary metabolites

Correlation analysis was conducted among the concentrations of individual secondary metabolites, antioxidant activities (ABTS and DPPH), TFC, and TPC in each species (Figure 5). The secondary metabolites used for the correlation analysis were those assigned to the highest statistical group for each species based on multiple comparison analysis. In soybean sprouts, significant correlations were observed among secondary metabolites, and these correlations were consistently positive. Daidzin showed moderate positive correlations with daidzein (r = 0.72), genistein (r = 0.76), and glycitin (r = 0.68). ABTS, TFC, and TPC showed positive correlations with biochanin A, daidzein, daidzin, genistein, and glycitin, with the exception of cinnamic acid. In particular, ABTS showed a statistically significant, very strong positive correlation with daidzein (r = 0.87) and daidzin (r = 0.85), as well as with TFC (r = 0.85), while statistically significant moderate positive correlations were observed with glycitin (r = 0.74) and TPC (r = 0.75). Additionally, cinnamic acid showed poor to fair negative association with daidzein (r = −0.19), daidzin (r = −0.49), genistein (r = −0.38), and glycitin (r = −0.34). In mungbean sprouts, a statistically significant positive correlation was observed between gallic acid and isoquercitrin (r = 0.66, moderate). In addition, a statistically significant positive correlation (r = 0.64, moderate) was observed between DPPH and TFC, while a statistically significant very strong positive correlation (r = 0.83) was observed between TPC and gallic acid. In cowpea sprouts, no statistically significant correlations were observed among species-specific secondary metabolites. However, daidzin exhibited statistically significant positive correlations with ABTS radical scavenging activity (r = 0.77), TFC (r = 0.53), and TPC (r = 0.64). The correlations between daidzin and ABTS, as well as between daidzin and TFC, were interpreted as fair, whereas the correlation between daidzin and TPC was interpreted as moderate. ABTS exhibited statistically significant positive correlations with TPC and TFC, with a particularly very strong positive correlation observed between ABTS and TPC (r = 0.95).

**Figure 5.**
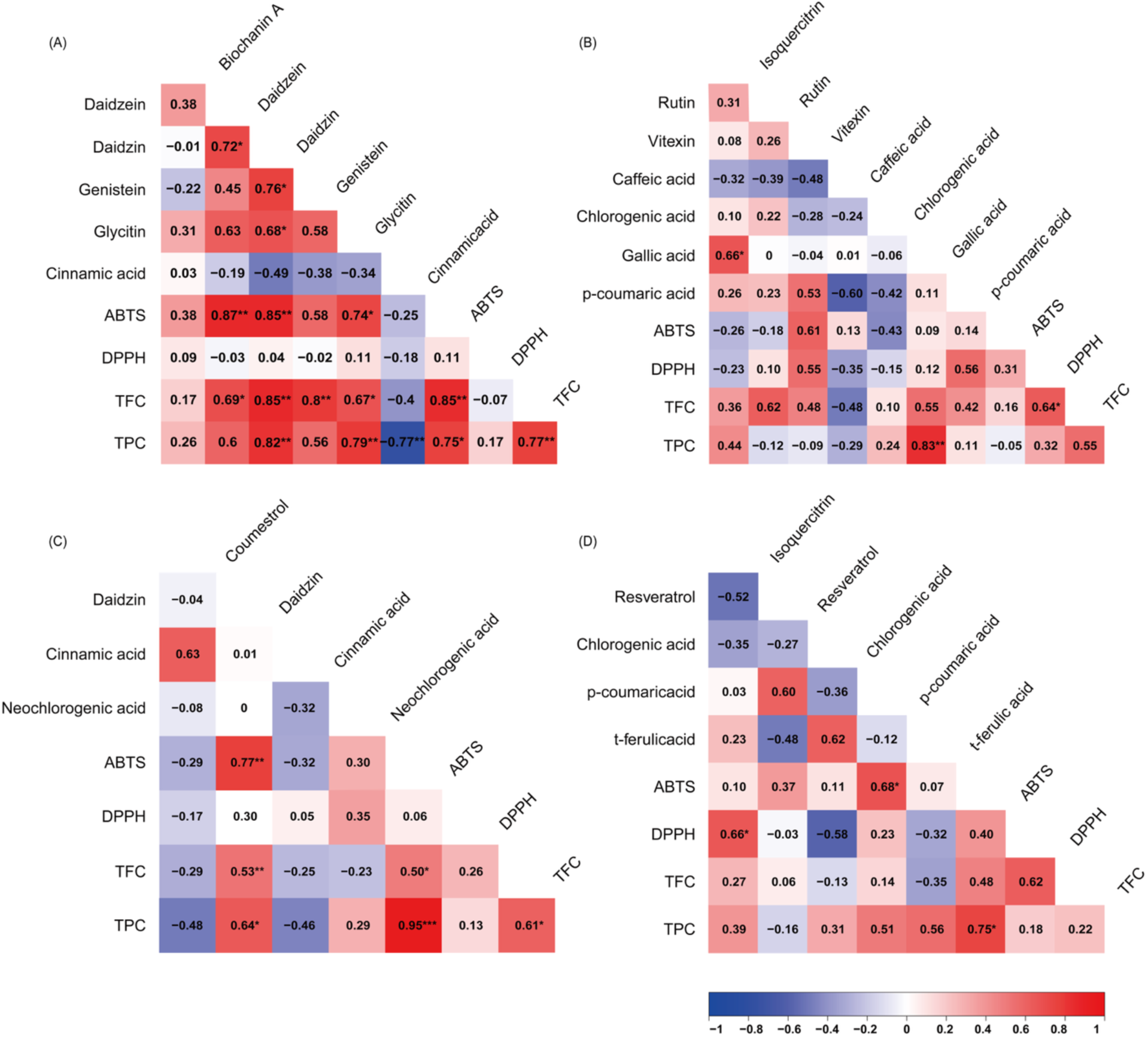
Correlation analysis between individual secondary metabolites, antioxidant activity, TFC, and TPC (A): soybean sprout, (B): mungbean sprout, (C): cowpea sprout, (D): peanut sprout Numbers shown in Figure 5 represent correlation coefficients, with asterisks (*, **, ***) indicating statistical significance (\**p* < 0.05, \*\**p* < 0.01, \*\*\**p* < 0.001).

Similarly, in peanut sprouts, no statistically significant correlations were detected among secondary metabolites. However, in peanut sprouts, statistically significant moderate positive correlations were observed between isoquercitrin and DPPH (r = 0.66), p-coumaric acid and ABTS (r = 0.68), and TPC and ABTS (r = 0.75).

## 4. Discussion

Germination is a method that can improve food quality in legume species by increasing antioxidant activity and the concentratons of specific secondary metabolites. The purpose of this study was to comparatively analyze antioxidant activity, TPC, total TFC, and individual secondary metabolite concentrations in soybean, mungbean, cowpea, and peanut sprouts. In this study, significant variability was observed among soybean, mungbean, cowpea, and peanut sprouts. In particular, mungbean sprouts showed significantly higher TPC, whereas mungbean and cowpea sprouts exhibited significantly higher TFC than the other three species (Figure 2). Consistently, mungbean and cowpea sprouts exhibited significantly higher antioxidant activity than soybean and peanut sprouts, supporting their potential as functional sprout vegetables. Additionally, statistically significant variability was also observed among accessions within species. In this study, distinct, species-specific secondary metabolite profiles were identified across the four legume sprouts (Figure 6). Previous studies have established that soybean is rich in isoflavones.^5^ Consistent with these reports, soybean sprouts in the present study contained higher levels of glycitin, daidzein, genistein, and biochanin A than mungbean, cowpea, and peanut sprouts. In mungbean sprouts, gallic acid, caffeic acid, vitexin, and rutin were present at higher concentrations than in the other species. Among these, rutin was particularly abundant, consistent with previous reports showing higher rutin levels in mungbean sprouts than in soybean and cowpea sprouts.^22^ Among these secondary metabolites, rutin exhibited the most pronounced species-dependent accumulation pattern, with concentrations in mungbean accessions generally exceeding those observed in accessions of the other three species. These findings suggest that rutin is a characteristic component of the secondary metabolite profile of mungbean sprouts among the legume sprouts examined in this study. Cowpea sprouts were characterized by significantly higher concentrations of neochlorogenic acid and coumestrol compared with the other species. Among these secondary metabolites, coumestrol is a phytoestrogen commonly found in animal feedstuffs and has been widely investigated in animal nutrition research.^23–24^ It occurs in the sprouts of various crops, including clover, alfalfa, and soybean.^23, 25–26^ Ohta *et al*. compared coumestrol contents in nine commercially available vegetables, including soybean sprouts, white radish sprouts, broccoli sprouts, pea sprouts, Chinese cabbage, Brussels sprouts, broccoli, cabbage, and soybeans, and reported that soybean sprouts contained the highest coumestrol concentration among the vegetables examined.^26^ However, the results of the present study demonstrated that cowpea sprouts exhibited significantly higher coumestrol concentrations than soybean sprouts, indicating that cowpea sprouts should also be considered when comparing coumestrol accumulation among legume sprouts. Although peanut sprouts exhibited relatively lower antioxidant activity, TFC, and TPC than the other legume sprouts, they showed a distinct species-biased metabolite profile. Chlorogenic acid, *t*-ferulic acid, resveratrol, and isoquercitrin were present at relatively higher levels in peanut sprouts than in the other three species. These findings suggest that peanut sprouts should be evaluated not only by overall indices such as antioxidant activity, TFC, and TPC, but also by their specific secondary metabolite composition. Taken together, these results demonstrate that each legume sprout species possesses a distinct secondary metabolite profile. These interspecific differences highlight the potential for selecting different legume sprouts according to specific nutritional or functional objectives. Furthermore, the considerable variation observed among accessions within each species suggests that accession selection should be considered alongside species selection.

**Figure 6.**
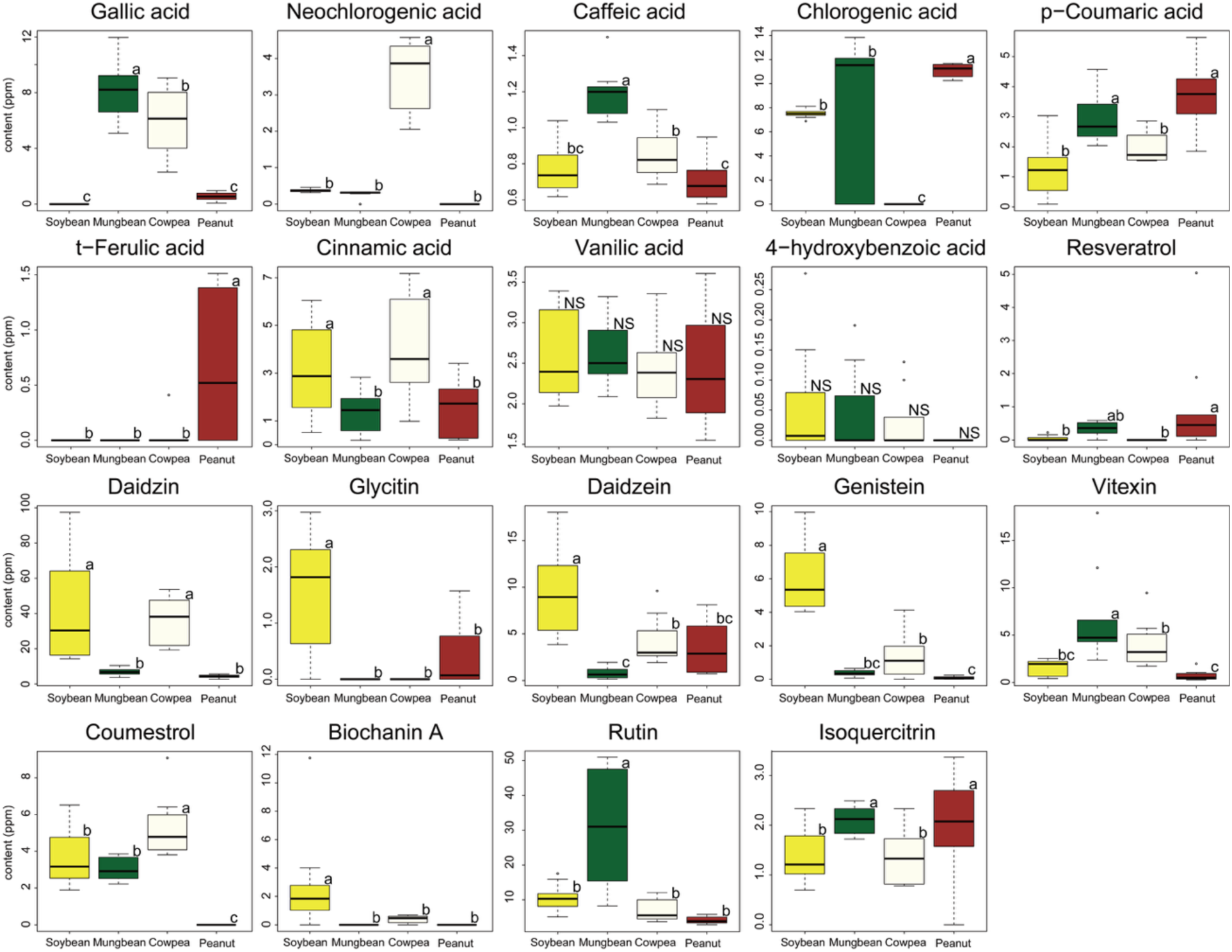
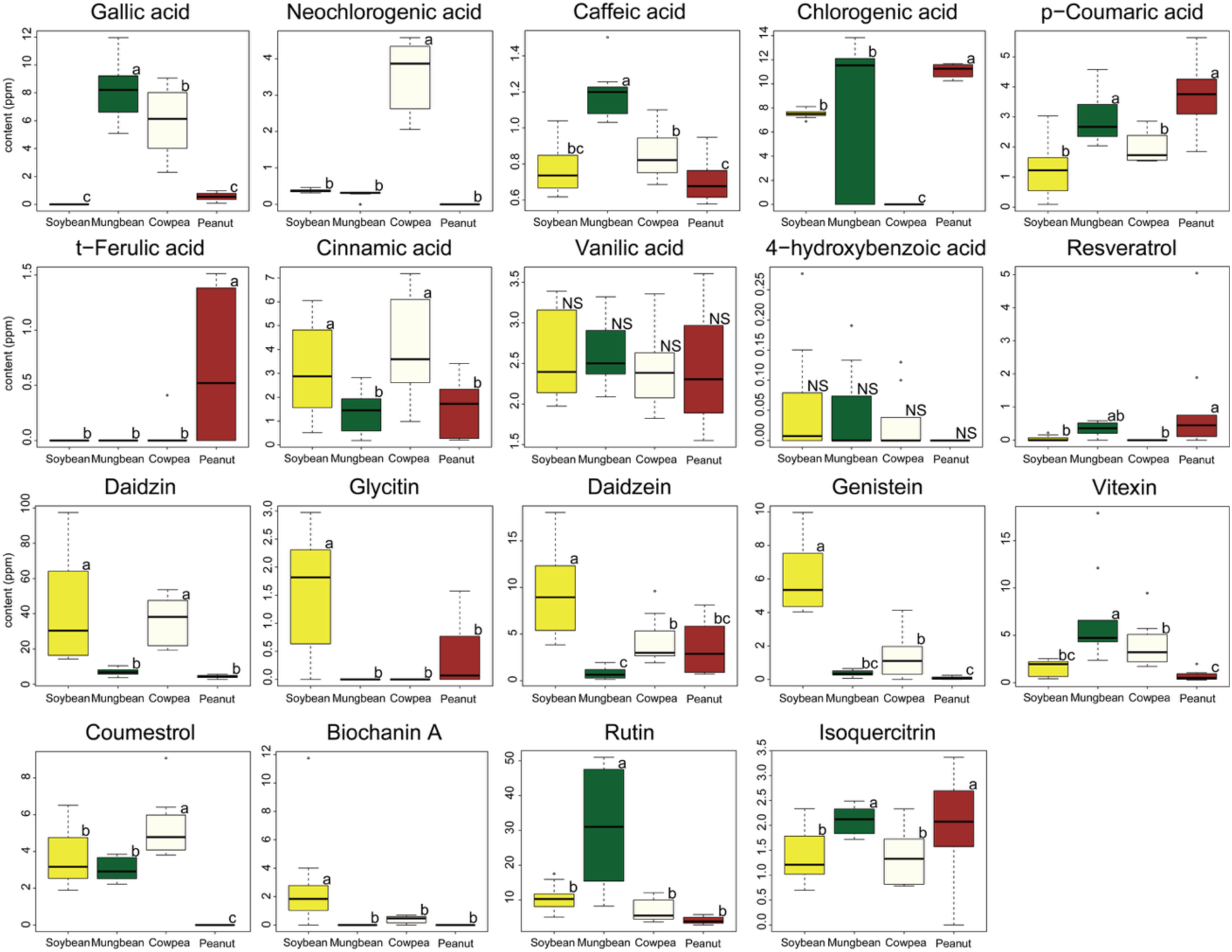
Comparison of individual secondary metabolites content between species of soybean, mungbean, cowpea, and peanut sprouts * NS: not significant Each lowercase letter is *p* < 0.05, indicating a significant difference between species.m

In general, antioxidant activity is associated with total phenolic content (TPC), and among phenols, flavonoids have been reported to be the major components with antioxidant activity in legumes.^27–28^ Therefore, correlation analysis was performed to evaluate the relationships among antioxidant activity, TPC, TFC, and representative secondary metabolites. Correlation analysis revealed diverse patterns in the magnitude and direction of the correlations among individual secondary metabolites, antioxidant activity, TFC, and TPC in each legume species (Figure 5). In soybean sprouts, cinnamic acid was negatively correlated with ABTS, DPPH, TFC, and TPC, whereas in cowpea sprouts, it showed a positive correlation with DPPH but negative correlations with ABTS, TFC, and TPC. Similarly, in mungbean sprouts, isoquercitrin showed a positive correlation with chlorogenic acid and negative correlations with ABTS and DPPH, whereas in peanut sprouts, the opposite pattern was observed. Overall, these differences in coefficient direction and magnitude suggest that the contribution of individual secondary metabolites to antioxidant activity and TFC and TPC of soybean, mungbean, cowpea, and peanut sprouts varies among species. Accordingly, antioxidant activity in legume sprouts cannot be fully elucidated by bulk indices such as TFC or TPC alone; rather, it appears to reflect the species-specific secondary metabolite composition and the distinct contributions of individual phenolic compounds.

In conclusion, this study demonstrated distinct interspecific and intraspecific variation in antioxidant-related traits and secondary metabolite profiles among legume sprouts, providing a comparative basis for selecting appropriate legume sprouts according to target nutritional or functional objectives.

## Abbreviations

ABTS: 2,2’-azino-bis(3-ethylbenzothiazoline-6-sulfonic acid)
DPPH: 2,2-diphenyl-1-picrylhydrazyl
TFC: Total flavonoid content
TPC: Total phenolic content
ANOVA: analysis of variance

## Funding

This work was supported by the New Faculty Startup Fund from Seoul National University.

## Supporting Information

List of accessions of soybean, mungbean, cowpea, and peanut sprouts (Table S1); List of analyzed secondary metabolites (Table S2);

## Notes

The authors declare no competing financial interest.

## ACKNOWLEDGMENT

The authors thank Prof. Bo-Keun Ha (Department of Applied Plant Science, Chonnam National University, Gwangju 61186, Republic of Korea) and Prof. Tae-Hwan Jun (Department of Plant Bioscience, Pusan National University, Miryang 50463, Republic of Korea) for providing the cowpea and peanut seeds, respectively, used in this study.

## Graphic for Table of Contents

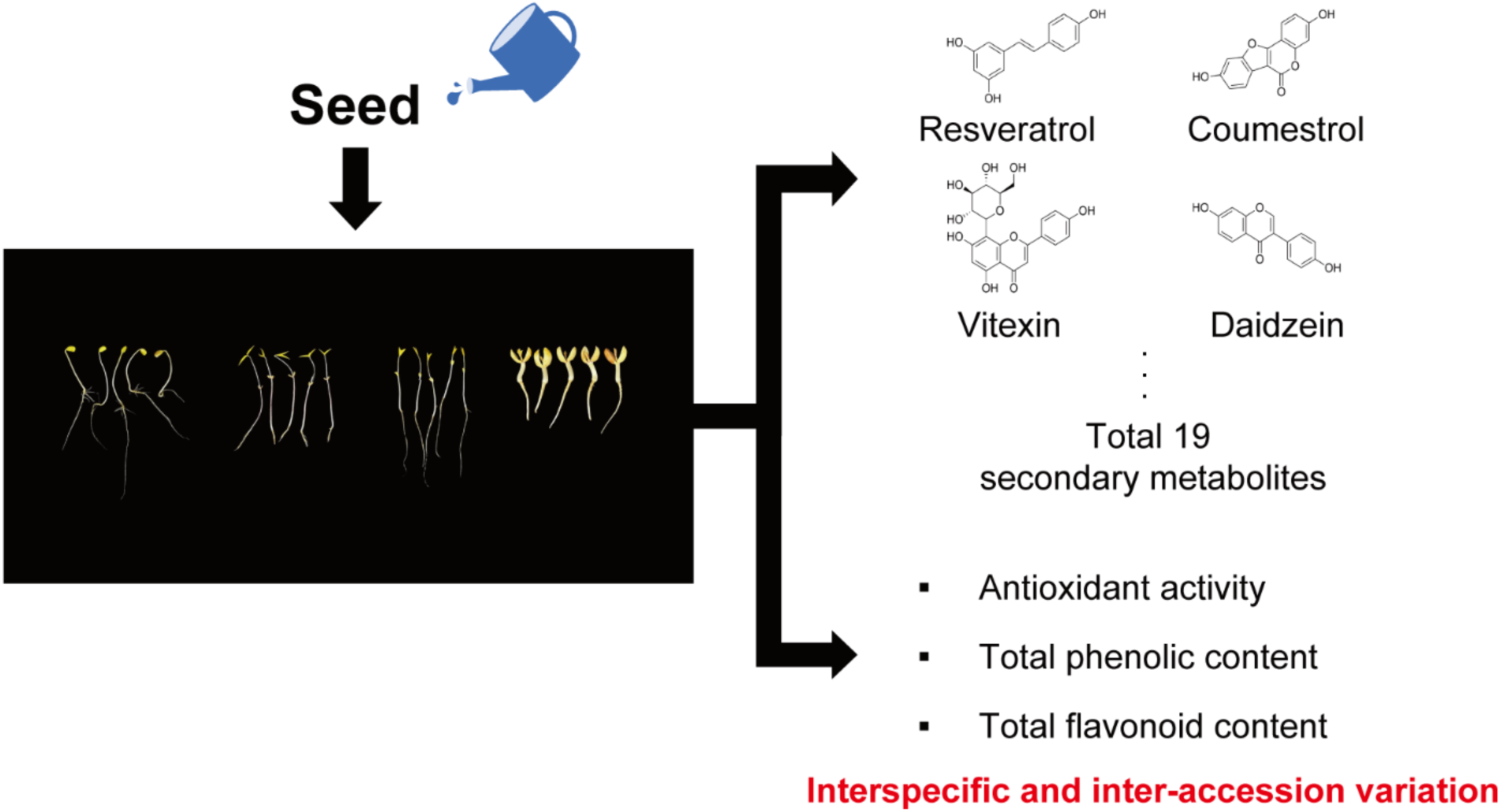

## Notes

### Competing Interest Statement

The authors have declared no competing interest.

