## supporting information for "Species and Accession Diversity of Secondary Metabolites and Antioxidant Activity in Legume Sprouts"

**Table S1.** List of soybean, mungbean, cowpea, and peanut accessions.

| Sample ID | | Accession | IT number | Origin | Sample ID | | IT number | Origin |
|---|---|---|---|---|---|---|---|---|
| Soybean | Cheongmiin | | IT 333602 | Korea | Cowpea | C81 | IT 105949 | Korea |
| | Chamame | | IT 185311 | Japan | | C93 | IT 111056 | Korea |
| | Daechan | | IT 333600 | Korea | | C123 | IT 145388 | Nigeria |
| | Seolitae | | IT 177849 | Korea | | C242 | IT 208369 | India |
| | Joyang | | IT 263170 | Korea | | C282 | IT 218033 | Thailand |
| | Pungsankong | | IT 263156 | Korea | | C292 | IT 228627 | Kyrgyzstan |
| | Haepum | | IT 270009 | Korea | | C319 | IT 237041 | Bulgaria |
| | SG151 | | IT 167957 | Bulgaria | | C365 | KHM-1 | Cambodia |
| | SG188 | | IT 213188 | Korea | | C369 | IT 712223 | Korea |
| | Gilyuk102 | | IT 323401 | Korea | | C378 | K016576 | Korea |
| Mungbean[16] | M658 | PB-1 (benggolo Puti) | | Indonesia | Peanut | P111 | PI 196635 | Madagascar |
| | Seonhwa | VC1973A | | Taiwan | | P189 | PI 269710 | Japan |
| | M366 | VO1946A-Y | IT 189516 | Philippines | | P310 | PI 403824 | Paraguay |
| | M334 | | IT162779 | Korea | | P366 | PI 494034 | Argentina |
| | M369 | V03720B-G | | USA | | P378 | PI 502116 | Peru |
| | Sanpo | | IT348878 | Korea | | P177 | PI 268885 | China |
| | M357 | | IT183264 | Korea | | P191 | PI 270783 | Zambia |
| | M243 | JP99066 | | Pakistan | | P277 | PI 355279 | Mexico |
| | M730 | W162 | | wild | | P294 | PI 381331 | Spain |
| | M625 | Arta ijo | | Indonesia | | P319 | PI 415870 | Senegal |

**Table S2**. Identified UPLC standard material details.

| Compound name | | Formula | Cas No. | RT | absorbance (nm) | $r^2$ |
|---|---|---|---|---|---|---|
| **Phenolic acids** | Gallic acid | $C_7H_6O_5$ | 149-91-7 | 2.91 | 280 | 0.99 |
| | Neochlorogenic acid | $C_{16}H_{18}O_9$ | 906-33-2 | 8.22 | 280 | |
| | 4-Hydroxybenzoic acid | $C_7H_6O_3$ | 99-96-7 | 10.18 | 260 | |
| | Caffeic acid | $C_9H_8O_4$ | 331-39-5 | 13.48 | 280 | |
| | Chlorogenic acid | $C_{16}H_{18}O_9$ | 327-97-9 | 13.97 | 280 | |
| | Vanillic acid | $C_8H_8O_4$ | 121-34-6 | 14.20 | 280 | |
| | p-Coumaric acid | $C_9H_8O_3$ | 501-98-4 | 20.97 | 280 | |
| | t-Ferulic acid | $C_{10}H_{10}O_4$ | 537-98-4 | 23.35 | 280 | |
| | Cinnamic acid | $C_9H_8O_2$ | 140-10-3 | 29.04 | 280 | |
| **Flavonoids** | Daidzin | $C_{21}H_{20}O_9$ | 552-66-9 | 22.64 | 280 | 0.99 |
| | Glycitin | $C_{22}H_{22}O_{10}$ | 40246-10-4 | 22.93 | 280 | |
| | Rutin | $C_{27}H_{30}O_{16}$ | 153-18-4 | 23.50 | 280 | |
| | Vitexin | $C_{21}H_{20}O_{10}$ | 3681-93-4 | 23.62 | 340 | |
| | Isoquercitrin | $C_{21}H_{20}O_{12}$ | 482-35-9 | 24.03 | 340 | |
| | Daidzein | $C_{15}H_{10}O_4$ | 486-66-8 | 27.91 | 280 | |
| | Genistein | $C_{15}H_{10}O_5$ | 446-72-0 | 29.98 | 280 | |
| | Coumestrol | $C_{15}H_8O_5$ | 479-13-0 | 30.12 | 240 | |
| | Biochanin A | $C_{16}H_{12}O_5$ | 491-80-5 | 35.00 | 280 | |
| **Stilbene** | Resveratrol | $C_{14}H_{12}O_3$ | 501-36-0 | 27.58 | 280 | 0.99 |

* RT = Retention Time (min)
